# Control of human testis-specific gene expression

**DOI:** 10.1101/593376

**Authors:** Jay C. Brown

## Abstract

**Background:** As a result of decades of effort by many investigators we now have an advanced level of understanding about several molecular systems involved in the control of gene expression. Examples include CpG islands, promoters, mRNA splicing and epigenetic signals. It is less clear, however, how such systems work together to integrate the functions of a living organism. Here I describe the results of a study to test the idea that a contribution might be made by focusing on genes specifically expressed in a particular tissue, the human testis.

**Experimental Design:** A database of 239 testis-specific genes was accumulated and each was examined for the presence of features relevant to control of gene expression. These include: (1) the presence of a promoter, (2) the presence of a CpG island (CGI) within the promoter, (3) the presence in the promoter of a transcription factor binding site near the transcription start site, (4) the level of gene expression, and (5) the above features in genes of cell types such as spermatocyte and spermatid that differ in their extent of differentiation.

**Results:** Of the 107 database genes with an annotated promoter, 56 were found to have one or more transcription factor binding sites near the transcription start site. Three of the binding sites observed, Pax-5, AP-2αA and GRα, stand out in abundance suggesting they may be involved in testis-specific gene expression. Compared to less differentiated testis-specific cells, genes of more differentiated cells were found to be (1) more likely to lack a CGI, (2) more likely to lack introns and (3) higher in expression level. The results suggest genes of more differentiated cells have a reduced need for CGI-based regulatory repression, reduced usage of gene splicing and a smaller set of expressed proteins

## Introduction

The regulatory control of gene expression is a central feature of all living organisms. Beginning with the same genome sequence, features of differential gene expression collaborate to create the entire landscape of tissue and cell function including a life-long developmental program, pathways to maintain homeostasis and functions able to respond to environmental change. The crucial importance of gene regulatory control has made it a thoroughly-studied and familiar area of investigation. As a result we now know about central features of regulation including the role of promoters, CpG islands, epigenetic signaling, transcription factors, enhancers, structured chromosome domains, mRNA splicing and many others [1-7]. Lacking, however, is an appreciation of how the individual systems work together to produce smoothly functioning developmental and other programs. Are there features that are more fundamental in that they are expressed earlier in development or affect a greater number of tissues and cells? To what extent is the pathway of gene regulatory systems the same in different tissues? Are there pathways of gene expression that use some but not all of the gene regulatory features used in others? Are regulatory features deployed differently in developmental pathways compared to those involved in response to environmental change? The above questions and many related ones currently occupy investigators studying gene regulatory control.

I have adopted the view that progress might be made by focusing on the genes specifically expressed in a single tissue. Limiting the analysis in this way significantly reduces the number of genes to be examined and also may reduce the number of regulatory systems that need to be considered. It is anticipated that information generated about regulation of genes expressed specifically in a single tissue may be able to be generalized to a larger and more diverse gene population.

Here I describe the results of studies carried out to examine genes expressed specifically in the human testis [8]. Testis is attractive for study because it consists predominantly of a highly restricted number (four) of distinct cell types that are all on the same pathway leading to production of a single cellular product, sperm [9]. Also, the testis stands out, compared to other tissues, for the high number of tissue-specific genes [10], a property that offers a similarly high number of regulatory features that might be relevant. Together the two features of testis, a small number of cells and a large number of specific genes, offer the possibility of relating control of specific gene expression to defined cellular developmental events.

The study began with creation of a database containing 239 genes expressed specifically in human testis. Database genes were chosen to be representative of the larger population of all testis-specific genes. The database includes genes encoded on all but one of the 24 human chromosomes; both protein-coding genes and genes that specify non-coding RNAs are represented. Database genes were examined for the presence and functioning of properties relevant to control of gene expression including the presence of a CpG island, the presence of a promoter, transcription factor binding sites within the promoter and the level of gene expression. The results are interpreted to clarify the role of the above features in control of testis-specific gene usage and their significance for sperm development.

## Materials and Methods

### Database of human testis-specific genes

The database of human testis-specific genes employed here (Table S1) contains 239 genes each annotated to be highly specific for testis in both the UCSC Genome Browser (version hg38, 2013 [https://genome.ucsc.edu/]) and the NCBI gene reference [https://www.ncbi.nlm.nih.gov/]. The database was curated from among genes contained in slightly larger databases of testis-specific genes [8, 11] and from a database of human gene promoters [12]. The goal was to create a gene set representative of all testis-specific genes.

### Gene properties examined

Genes with a CpG island (CGI) were identified from the UCSC Genome Browser (version hg38, 2013). All database testis-specific genes with an annotated CGI near the transcription start site (TSS) were included without regard to the length of the CGI or its percent GC content. Genes containing a promoter were identified by the FirstEF algorithm [13] as found in the 2003 (hg36) version of the UCSC Genome Browser. For all genes examined, the level of testis-specific expression was retrieved from the UCSC Genome Browser (version hg38, 2013). A gene was considered to be broadly expressed if it was annotated to have a comparable level of expression in half or more of the tissues reported in the UCSC or NCBI databases. The list of Djureinovic et al. [8] was used to identify gene-encoded proteins highly enriched in spermatogonia, spermatocyte, spermatid or sperm. Genes lacking introns were identified using the Intronless Gene Database (http://www.bioinfo-cbs.org/igd/).

### Transcription factor binding sites

Transcription factor binding sites (TFBS) near transcription start sites were identified beginning with promoters downloaded from the UCSC Genome Browser [12]. Promoters were identified by the FirstEF algorithm as described above. Each was 1000bp in length beginning 570bp upstream from the TSS and ending 430bp downstream. The entire 1000bp promoter sequence was scanned for the presence of TFBS with the ALGGEN-PROMO website running TRANSFAC version 8.3 (maximum matrix dissimilarity rate=2; http://alggen.lsi.upc.es/cgi-bin/promo_v3/promo/promoinit.cgi?dirDB=TF_8.3). TFBS or combinations of contiguous TFBS were included in Table S1 if they were found between - 10bp and +10bp of the annotated TSS and were 6bp or more in length.

## Results

### Testis-specific gene database

Database testis-specific genes were found to be widely distributed among the 24 human chromosomes. All but the Y chromosome encode at least one testis-specific database gene. Chromosome 1 has the most (28 of 239 database genes) and chromosome 21 the least (1 gene; Fig. 1a). When expressed as the number of database genes per 100Mb of chromosome sequence, the highest number was found in chromosome 19 (19.0) and the lowest in chromosome 21 (2.2; Fig. 1b).

**Fig 1:**
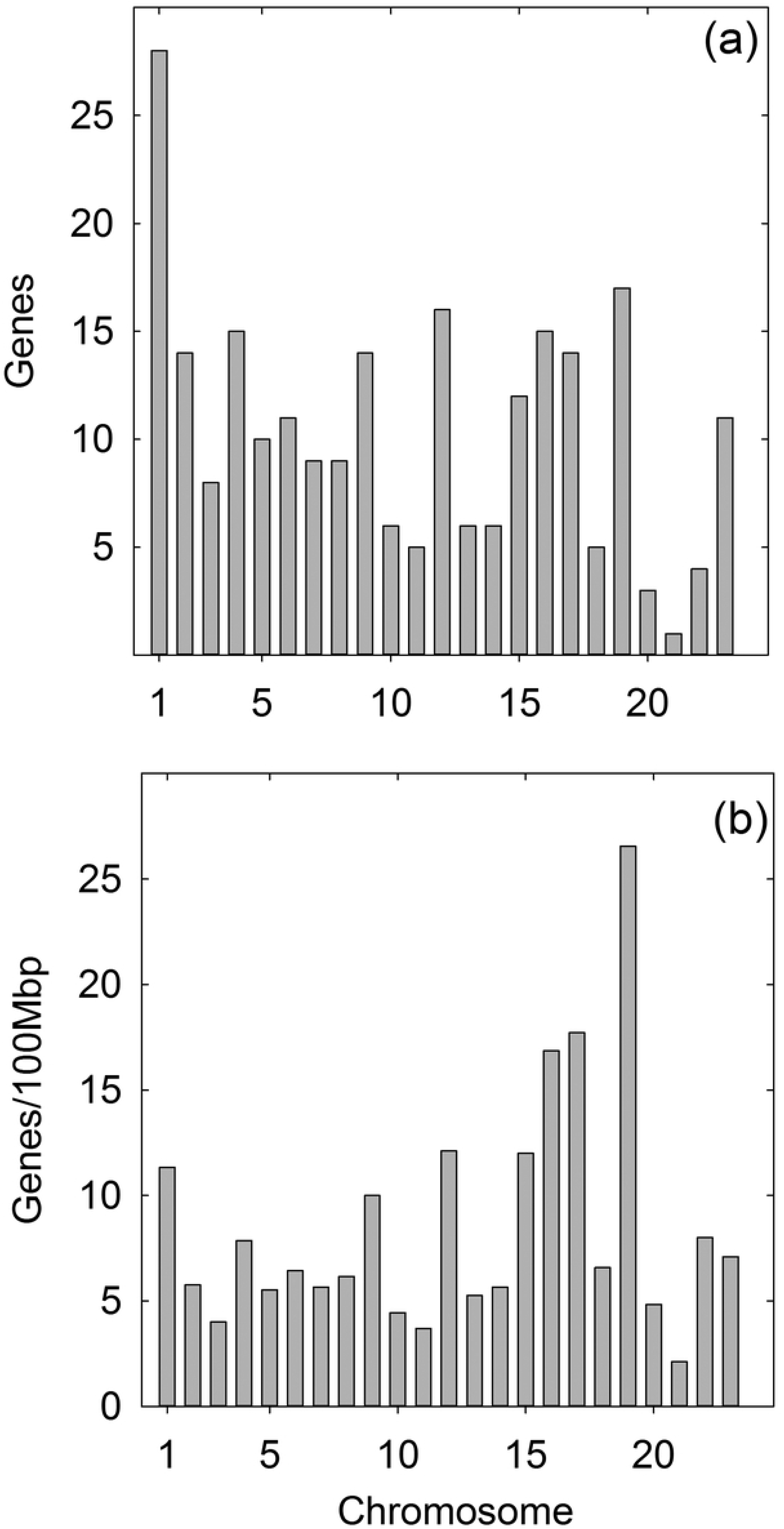
Chromosome distribution of database human testis-specific genes. (a) Number of database genes encoded on each chromosome. (b) Gene density expressed as genes/100mbp of chromosome sequence. Note the high density of database testis-specific genes on chromosome 19.

The expression level of database genes was found to favor those with low expression. For instance, 194 of the 239 genes (81%) have expression levels in the lowest 1/3 of the distribution (Fig. 2). Among the highly expressed genes, the distribution shows preferred values of ~60, 170 and 215 RPKM suggesting there may be a mechanism to favor particular expression levels (Fig. 2).

**Fig 2:**
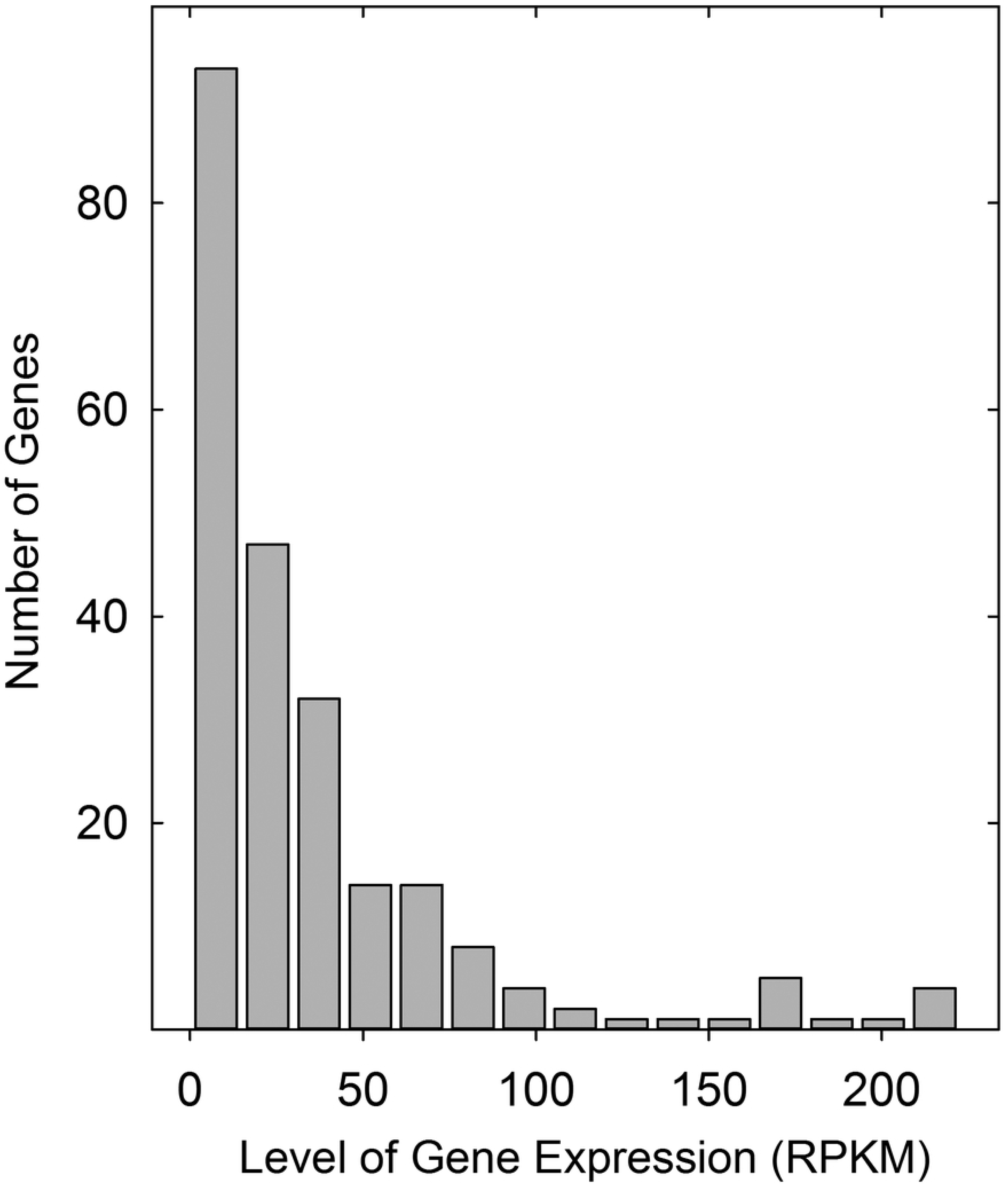
Histogram showing the expression level of all database testis-specific human genes. Note that expression level is skewed to the low expression end of the distribution.

### CpG islands in testis-specific human genes

As tissue specific genes have been reported to be depleted in CpG islands compared to broadly expressed genes [2, 14, 15], it was expected that database testis-specific genes would be depleted in CGI, and this was found to be the case (Table 1). Of the 239 database genes, 127 (53.1%) were found to lack a CGI. In contrast, absence of a CGI was observed in only 8.0% of an unselected human gene population and 9.4% of a population of broadly expressed genes (Table 1). Testis-specific LINC genes were almost all lacking a CGI (14 of 15 LINC genes) while among testis-specific intronless genes the proportion was about the same as the testis-specific population as a whole (50.0% for intronless genes compared to 53.1% for all database genes; Table 1).

**Table 1:** Testis-specific genes lacking a CpG island

| Gene Population | Genes lacking a CpG Island<sup>a</sup> |
| --- | --- |
| Testis-specific (all database) | 127/239 53.1% |
| Unselected human genes<sup>b</sup> | 8/100 8.0% |
| Broadly-expressed human genes<sup>c</sup> | 10/106 9.4% |
| Testis-specific LINC genes | 14/15 93.3% |
| Testis-specific intronless genes | 12/24 50.0% |

<sup>a</sup> Data from UCSC Genome Browser Reference Human Genome version hg38, 2013.
<sup>b</sup> Sequential genes on chromosome 9 beginning with ACO1.
<sup>c</sup> Sequential broadly-expressed genes on chromosome 12 beginning with ZNF641.

Testis-specific database genes lacking a CGI did not differ greatly in expression level from CGI-containing testis-specific genes or from all testis-specific genes; mean expression levels were 48.4, 50.7 and 49.5 RPKM, respectively (Table 2). This result suggests CGI are not directly involved in determining gene expression level. The observation is compatible with the accepted view that CGI function in large-scale gene repression by way of methylation, a modification that suppresses expression of affected genes [2, 16].

**Table 2:**
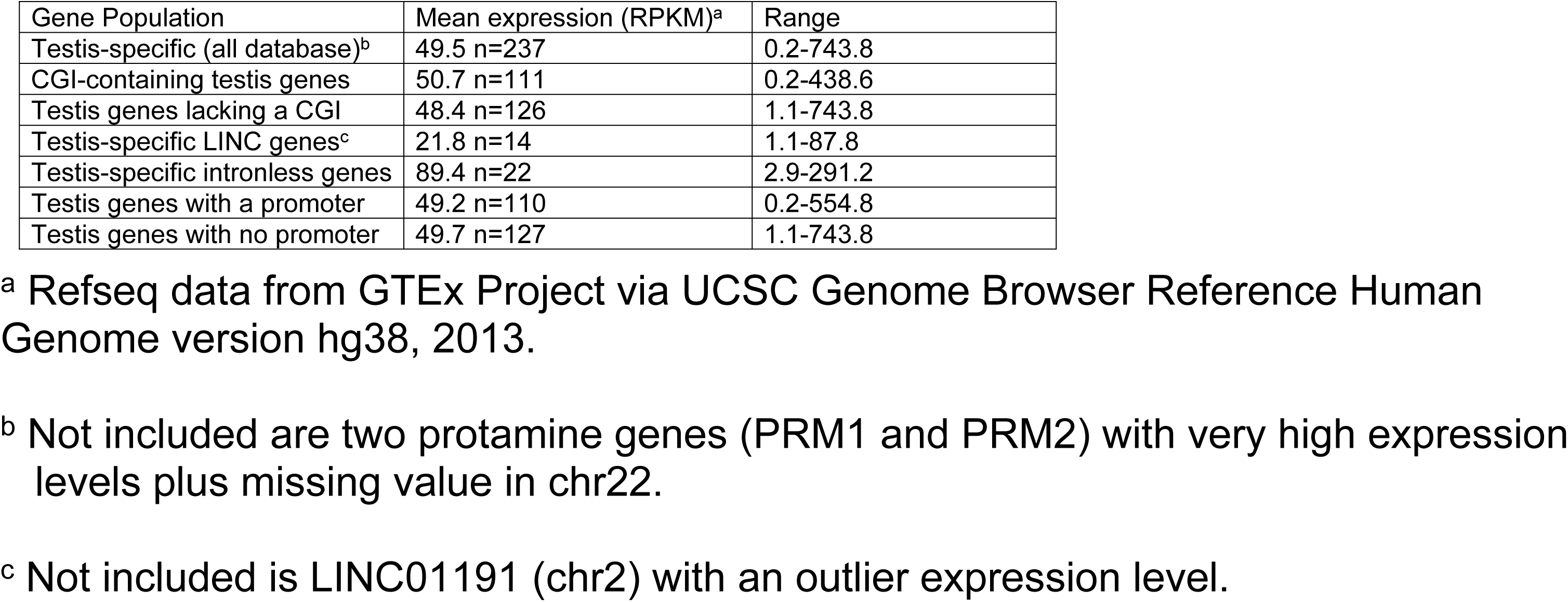
Expression level of testis-specific gene populations

| Gene Population | Mean expression (RPKM)<sup>a</sup> | Range |
| --- | --- | --- |
| Testis-specific (all database)<sup>b</sup> | 49.5 n=237 | 0.2-743.8 |
| CGI-containing testis genes | 50.7 n=111 | 0.2-438.6 |
| Testis genes lacking a CGI | 48.4 n=126 | 1.1-743.8 |
| Testis-specific LINC genes<sup>c</sup> | 21.8 n=14 | 1.1-87.8 |
| Testis-specific intronless genes | 89.4 n=22 | 2.9-291.2 |
| Testis genes with a promoter | 49.2 n=110 | 0.2-554.8 |
| Testis genes with no promoter | 49.7 n=127 | 1.1-743.8 |

<sup>a</sup> Refseq data from GTEx Project via UCSC Genome Browser Reference Human Genome version hg38, 2013.
<sup>b</sup> Not included are two protamine genes (PRM1 and PRM2) with very high expression levels plus missing value in chr22.
<sup>c</sup> Not included is LINC01191 (chr2) with an outlier expression level.

### Expression levels of testis-specific LINC and intronless genes

Testis-specific long, intergenic non-coding (LINC) genes were found to have a lower mean expression level compared to all testis-specific database genes. The difference was ~2.2 fold (49.5 RPKM compared to 21.8; Table 2). This observation is in qualitative agreement with results showing decreased expression of LINC genes in databases of all human LINC genes [17, 18]. In contrast, database testis-specific intronless genes were found to have a mean expression level higher than that of all testis-specific genes (89.4RPKM compared to 49.5; Table 2). This observation indicates that testis-specific intronless genes must possess strong nuclear export and other translation-enabling features that do not depend on the presence of introns and mRNA splicing pathways [19, 20].

### Testis-specific genes with a promoter

As promoters can play an important role in control of gene expression, they were examined carefully in the testis-specific population considered here. Special attention was devoted to transcription factor binding sites (TFBS) near the annotated transcription start site because such TFBS can have a direct effect on initiation of new gene transcription [21, 22]. Less than half of the database testis-specific genes were found to have an annotated promoter (107/239 genes; 44.8%; see Table 3). This compares to greater than 90% in a population of unselected human genes. Both LINC and intronless testis-specific gene populations were also found to be depleted in promoter-containing genes. Percentages were 6.7% (1/15) of LINC genes with a promoter and 29.1% (7/24) for intronless genes (Table 3). The lower number of promoter-containing genes in the testis-specific population suggests that in many testis-specific genes the functions of the promoter must be accomplished by unannotated promoters or by other gene features.

**Table 3:** Testis-specific genes with a promoter and transcription factor binding site near the transcription start site

| Gene Population | Genes with promoter<sup>a</sup> | Promoters with TFBS at TSS<sup>b</sup> |
| --- | --- | --- |
| Testis-specific (all database) | 107/239 44.8% | 56/107 52.3% |
| Unselected human genes<sup>c</sup> | 94/104 90.4% | 51/141 36.2% |
| Testis-specific LINC genes | 1/15 6.7% | 1/1 100.0% |
| Testis-specific intronless genes | 7/24 29.1% | 3/7 42.9% |

<sup>a</sup> Data from UCSC Genome Browser Reference Human Genome version hg38, 2013.
<sup>b</sup> Within 10bp of the transcription start site.
<sup>c</sup> Sequential genes on chromosome 9 beginning with ACO1.

The ALGGEN-PROMO web site was used to retrieve transcription factor binding sites near the TSS in database gene promoters as described in Materials and Methods. A total of 25 different transcription factor binding sites were observed among the 56 genes with a promoter (see Tables S2 and 3). Highest in abundance were Pax-5, AP-2αA and GR-α which were present in 12, 10 and 8 gene promoters, respectively (Table 4). Together the three account for 30 of the 56 transcription factor binding sites (53.5%) present in relevant database genes suggesting they may have a role in regulation of testis-specific gene expression. Eleven of the 25 different transcription factor binding sites were each present near the TSS in only one database gene promoter (Table S2).

**Table 4:** Transcription factor recognition sequence near TSS in the promoters of database human testis-specific genes

| Transcription Factor<sup>a</sup> | Sequence recognized | No. genes | Mean. Expression (RPKM) | Range (RPKM) |
| --- | --- | --- | --- | --- |
| Pax-5 | GCCC | 12 | 44.3 | 12.5-174.2 |
| AP-2αA | GCAGGC | 10 | 80.6 | 5.2-422.7 |
| GR-α | ANAGGGR<sup>b</sup> | 6 | 132.4 | 2.2-554.8 |
| GR-α | CCTCT | 2 | 34.4 | 21.5-78.1 |

<sup>a</sup> Most abundant three transcription factors binding sites near TSS among 25 TFBS in 56 testis-specific genes.
<sup>b</sup> N= A, T, G or C; R= A or G

The nucleotide sequences of transcription factor binding sites were also retrieved in case they might suggest the identity of other elements that recognize the same DNA sites (Table S2). The sequences were scanned visually to identify similarities, and the results are summarized in Table 4. One recurring site was found to correspond to the Pax-5 binding site, one for AP-2αA and two for GR-α (Table 4). The four sequences suggest themselves as candidates for a role in control of testis-specific gene expression. In each sequence the relevant genes were found to vary significantly in level of expression indicating that the sequences and the transcription factors that bind them may act to activate or repress gene expression depending on the context of other regulatory features present (Tables S2 and 4).

### Sperm progenitor cells in the testis

Seminiferous tubules are the major structural feature of the testis accounting for more than 80% of the testis mass. They consist of six distinct cell types. Four are direct precursors of sperm (the spermatogonia, spermatocytes, spermatids and sperm themselves), while two others support spermatogenesis but do not themselves develop into sperm (Leydig and Sertoli cells). Sperm progenitor cells are arranged radially in the seminiferous tubule with the spermatogonia located furthest from the tubule lumen and spermatocytes, spermatids and sperm progressively nearer [23, 24].

Spermatogonial cells divide to produce: (1) primary spermatocytes capable of further differentiation to create sperm; and (2) cells capable of replenishing the spermatogonial population. Both spermatogonium progeny cell types are diploid. Primary spermatocytes undergo a meiotic division to produce secondary spermatocytes. These are haploid cells that divide to produce spermatids, cells that further differentiate to become sperm. The well-characterized pathway leading to sperm production described above creates an opportunity to ask how features controlling gene expression may correlate with and underlie the molecular events involved. Below I describe studies designed to clarify how aspects of gene regulatory control may be involved.

The studies were enabled by the existence of a database of 122 testis-specific genes whose expression has been defined in individual sperm pathway cell types [8]. Assignments were made by noting the binding of protein-specific antibodies to sections of seminiferous tubule tissue. If the cell type(s) was defined for a testis-specific gene examined here, it is noted in Table S1. The results showed that the cell type(s) was defined for 40 of the 239 database genes. As shown in Table 5, four database genes were found in spermatogonium, 7 in spermatocytes, 17 in spermatid and 17 in sperm. Features of gene regulatory control noted were: (1) the presence of a CGI in the promoter, (2) the presence of introns in the gene and (3) the gene expression level (Table 5).

**Table 5:** Gene expression in sperm lineage cells

| Cell type | Database genes with |  | Mean expression (RPKM)<sup>a</sup> | Range (RPKM) |
| --- | --- | --- | --- | --- |
|  | No CG island | No Introns |  |  |
| Spermatogonium | 1/4 (25%) | 1/5 (20%) | 24.1 n=6 | 7.7-58.7 |
| Spermatocyte | 0/7 (0%) | 1/8 (12%) | 42.3 n=8 | 7.7-93.6 |
| Spermatid | 9/17 (53%) | 4/16 (25%) | 91.9 n=13 | 29.5-255.1 |
| Sperm<sup>b</sup> | 11/17 (65%) | 6/18 (33%) | 101.1 n=13 | 9.2-255.1 |

<sup>a</sup> Refseq data from GTEx Project via UCSC Genome Browser Reference Human Genome version hg38, 2013.
<sup>b</sup> Not included are two protamine genes (PRM1 and PRM2) with very high expression levels

The results in the case of CG islands show that the population of genes expressed at early stages of sperm formation (spermatogonia and spermatocytes) has a lower proportion of CGI-negative genes compared to the more differentiated cells (i.e. spermatid and sperm). The proportion in less differentiated cells more closely resembles that seen in unselected human gene populations (8.0%; see Table 1) than in all testis-specific genes (53.1%). In the more differentiated cells, however, the proportion is more similar to the population of all testis-specific genes (i.e. 53% and 65% compared to 53.1%). The result suggests that more differentiated cells are better able to function without a CGI or do not have a need for a CGI. This would be the case, for instance, if more differentiated cells do not require large scale, more permanent gene repression by the CGI methylation pathway.

The proportion of intronless genes in sperm precursor cell types was found to be lower than in the proportion in all testis-specific genes (i.e. 12%-33% compared to 50%; see Tables 5 and 1). If this result is not affected by the small number of pathway-specific genes available for analysis, then it indicates that sperm precursor cells may benefit from gene splicing and nuclear export events found in splicing pathways.

Finally, the expression level of genes in less differentiated cell types was found to be lower than those in more highly differentiated ones (Table 5). Levels in less differentiated cells were lower than the average for all testis-specific genes (i.e. 24.1 and 42.3 compared to 49.4 RPKM) while higher levels were observed in the more differentiated populations. This observation is consistent with the idea that as cells differentiate they express a smaller number of distinct genes, but genes in the group are expressed at a higher level.

## Discussion

### Control of gene expression

Current ideas about vertebrate gene regulation emphasize the involvement of structured chromosomal domains [25-27]. Actively expressed genes are thought to be contained on regions of chromatin that project outward from a core region of heterochromatin, an area where gene expression is repressed. Projecting or looped chromatin regions contain a small number of active genes located between insulator regions composed of CTCF/ cohesion or YY1 binding sites [26, 28]. Active genes present in loops contain RNA polymerase II (RNAPII), promoters and transcription factors involved in gene regulatory control. Also present may be enhancer/promoter regions of DNA located remotely on the chromosome, but containing bound transcription factors able to affect gene expression.

### CG islands

CGI-containing genes suggest themselves as components of the heterochromatic region where gene expression is suppressed. Methylation of CpG sequences is known to repress gene expression or to make temporary repression more permanent [2]. The absence of CGI from a substantial portion (~50%; Table 1) of the testis-specific genes examined here suggests CGI may be a threat to testis-specific gene expression if genes were able to be suppressed by CpG methylation. If repression is appropriate, then it might be safer to do so by a more targeted, less permanent mechanism. In contrast to the testis-specific genes that lack a CGI, the results here show that a significant proportion has a CGI (also ~50%; Table 1). This would be the case with genes whose expression needs to be suppressed in non-testis tissues.

LINC genes constitute a second population where many genes lack CGIs (Table 1). LINC are weakly expressed genes that specify non-coding RNA molecules thought to function as sponges for unneeded proteins or perhaps as components of protein-RNA complexes [17, 29]. The lack of CGIs in most LINC gene promoters suggests it is rarely necessary for their expression to be repressed permanently.

### Level of gene expression

Current ideas about the role of structured chromosome domains provide few clues regarding factors that affect the level of gene expression. Proximity of a gene to a CTCF/cohesion insulator may potentiate expression, but otherwise little guidance is provided [26]. The results reported here indicate that a gene’s expression level is not strongly affected by a CGI in the promoter region. The mean expression level of genes with a CGI in the promoter is about the same as that of genes lacking a CGI (Table 2). Also, LINC genes were found to be more weakly expressed compared to the average of testis-specific genes, and intronless genes are more strongly expressed. The latter observation is in conflict with results indicating that the level of gene expression is potentiated by the presence of introns and mRNA splicing pathways [30, 31].

### Transcription factor binding

As it is well established that transcription factors bound to the promoter can have important effects on gene expression, transcription factor binding sites were examined thoroughly in the testis-specific gene population considered here. To simplify the analysis somewhat, I focused only on TFBS near the transcription start site. This simplification can be justified by the fact that the TSS is the site where transcription by RNAPII is initiated and where binding of a transcription factor might have its maximum effect.

The results led to the identification of three transcription factors (Pax-5, AP-2αA and GR-α) whose abundance make them candidates for a role in testis-specific gene expression (Table 4). Although Pax-5 is best known for its effects on B cell development, it has been noted to be prominently expressed in testis [32 33]. A similar situation applies in the case of AP-2αA. While AP-2α is best known for effects in the nervous system [34, 35], a related transcription factor, AP-2γ, recognizes a DNA sequence similar to that of AP-2αA and has effects on testis development [36, 37]. I suggest that AP-2γ could be the factor that recognizes AP-2 sites in the testis-specific genes identified here. GRα, a member of the glucocorticoid receptor family, is widely expressed in human tissues where it is known to have multiple effects on gene expression [38]. It would have specific effects in the testis only if another feature such as a specific isoform or association with another protein were involved [39].

As shown in Table 4, a wide range of expression level was observed among the genes having a TSS-proximal TF. For instance in the case of genes having a Pax-5 TF site, the range was 12.5-174.2 RPKM. This observation suggests the effect of individual TFs can be either activating or suppressive.

### Differentiation of testis-specific cells

The present study benefitted from the results of immuno-histochemical analyses in which testis-specific genes could be associated with cells at progressively more mature states of differentiation [8]. All four recognized pathway-specific cell types were found to be populated by at least a few database testis-specific genes (Table 5). This permitted features of gene regulatory control to be compared among genes of the four cell types (i.e. spermatogonium, spermatocyte, spermatid and sperm). The results showed that an increase in differentiated state correlated with an increase in the proportion of genes: (1) lacking a CGI, (2) lacking introns, and (3) with an increased level of gene transcription. The observed increase in the proportion of genes lacking a CGI may be interpreted in the same way as the similar increase observed in the case of broadly expressed compared to tissue specific genes [2, 15]. Genes of more highly specialized cells (i.e. tissue specific and more differentiated cells) may have a reduced need for permanent repression by the CpG methylation pathway. A similar interpretation is suggested to apply to the observed increase in intronless genes among more highly differentiated cells. Such genes may be reduced in their need for mRNA splicing and splicing-related pathways of mRNA transport out of the nucleus. The observed increase in tissue-specific gene expression level with cell differentiation state (Table 5) may be simply a consequence of the overall differentiation process. As a more highly specialized cell is created, the need for more abundant, highly specialized gene products is increased while products of less specialized cells is decreased.

The observed increase in expression level in more differentiated cells could have a useful consequence for investigators studying gene regulation. The correlation of expression level with increased differentiation state could be used to identify the extent of differentiation in an unknown cell type.

Finally, focus on a population of tissue specific genes as described here is interpreted to support the view that this is an attractive way to further our understanding of development and cell differentiation processes. It might be of interest, for instance, to know whether the observed increase in CGI-less genes and intronless genes observed here with more differentiated testis-specific genes is also found in specific genes of other tissues. Additional features of gene regulatory control such as the role of insulators and structural domains might also be productively evaluated with tissue-specific genes.

### Supporting Information

Table S1: Database of human testis-specific genes

Table S2: Human database testis-specific genes with a promoter and a transcription factor binding site near the transcription start site

## Acknowledgments

I thank Forde Upshur and Karsten Siller for invaluable programming support for this project.

## Author Contributions

All contributions: Jay C. Brown

## References

1. Gagniuc P, Ionescu-Tirgoviste C. Eukaryotic genomes may exhibit up to 10 generic classes of gene promoters. BMC Genomics. 2012;13:512.

2. Deaton AM, Bird A. CpG islands and the regulation of transcription. Genes Dev. 2011;25(10):1010–22.

3. Portela A, Esteller M. Epigenetic modifications and human disease. Nat Biotechnol. 2010;28(10):1057–68.

4. Lee TI, Young RA. Transcription of eukaryotic protein-coding genes. Annu Rev Genet. 2000;34:77–137.

5. Hnisz D, Abraham BJ, Lee TI, Lau A, Saint-Andre V, Sigova AA, et al. Superenhancers in the control of cell identity and disease. Cell. 2013;155(4):934–47.

6. Dixon JR, Selvaraj S, Yue F, Kim A, Li Y, Shen Y, et al. Topological domains in mammalian genomes identified by analysis of chromatin interactions. Nature. 2012;485(7398):376–80.

7. Black DL. Mechanisms of alternative pre-messenger RNA splicing. Annu Rev Biochem. 2003;72:291–336.

8. Djureinovic D, Fagerberg L, Hallstrom B, Danielsson A, Lindskog C, Uhlen M, et al. The human testis-specific proteome defined by transcriptomics and antibody-based profiling. Mol Hum Reprod. 2014;20(6):476–88.

9. Trainer TD. Histology of the normal testis. Am J Surg Pathol. 1987;11(10):797–809.

10. Uhlen M, Fagerberg L, Hallstrom BM, Lindskog C, Oksvold P, Mardinoglu A, et al. Proteomics. Tissue-based map of the human proteome. Science 2015;347(6220):1260419.

11. Liu Y, Jiang M, Li C, Yang P, Sun H, Tao D, et al. Human t-complex protein 11 (TCP11), a testis-specific gene product, is a potential determinant of the sperm morphology. Tohoku J Exp Med. 2011;224(2):111–7.

12. Brown JC. Control of human gene expression: High abundance of divergent transcription in genes containing both INR and BRE elements in the core promoter. PLoS One. 2018;13(8):e0202927.

13. Davuluri RV, Grosse I, Zhang MQ. Computational identification of promoters and first exons in the human genome. Nat Genet. 2001;29(4):412–7.

14. Vinson C, Chatterjee R. CG methylation. Epigenomics. 2012;4(6):655–63.

15. Zhu J, He F, Hu S, Yu J. On the nature of human housekeeping genes. Trends Genet. 2008;24(10):481–4.

16. Bogdanovic O, Veenstra GJ. DNA methylation and methyl-CpG binding proteins: developmental requirements and function. Chromosoma. 2009;118(5):549–65.

17. Cabili MN, Trapnell C, Goff L, Koziol M, Tazon-Vega B, Regev A, et al. Integrative annotation of human large intergenic noncoding RNAs reveals global properties and specific subclasses. Genes Dev. 2011;25(18):1915–27.

18. Derrien T, Johnson R, Bussotti G, Tanzer A, Djebali S, Tilgner H, et al. The GENCODE v7 catalog of human long noncoding RNAs: analysis of their gene structure, evolution, and expression. Genome Res. 2012;22(9):1775–89.

19. Grzybowska EA. Human intronless genes: functional groups, associated diseases, evolution, and mRNA processing in absence of splicing. Biochem Biophys Res Commun. 2012;424(1):1–6.

20. Lei H, Dias AP, Reed R. Export and stability of naturally intronless mRNAs require specific coding region sequences and the TREX mRNA export complex. Proc Natl Acad Sci U S A. 2011;108(44):17985–90.

21. Smale ST, Kadonaga JT. The RNA polymerase II core promoter. Annu Rev Biochem. 2003;72:449–79.

22. Roy AL, Singer DS. Core promoters in transcription: old problem, new insights. Trends Biochem Sci. 2015;40(3):165–71.

23. Ross MH, Pawlina W. Histology : a text and atlas : with correlated cell and molecular biology. 5th ed. Baltimore, MD: Lippincott Wiliams & Wilkins; 2006. xvii, 906 p. p.

24. Hill, M.A. (2019, March 18) Embryology Spermatozoa Development. Retrieved from https://embryology.med.unsw.edu.au/embryology/index.php/Spermatozoa_Development.

25. Li G, Ruan X, Auerbach RK, Sandhu KS, Zheng M, Wang P, et al. Extensive promoter-centered chromatin interactions provide a topological basis for transcription regulation. Cell. 2012;148(1-2):84–98.

26. Tang Z, Luo OJ, Li X, Zheng M, Zhu JJ, Szalaj P, et al. CTCF-Mediated Human 3D Genome Architecture Reveals Chromatin Topology for Transcription. Cell. 2015; 163(7):1611–27.

27. van Steensel B, Belmont AS. Lamina-Associated Domains: Links with Chromosome Architecture, Heterochromatin, and Gene Repression. Cell. 2017;169(5):780–91.

28. Weintraub AS, Li CH, Zamudio AV, Sigova AA, Hannett NM, Day DS, et al. YY1 Is a Structural Regulator of Enhancer-Promoter Loops. Cell. 2017;171(7):1573–88 e28.

29. Ulitsky I, Bartel DP. lincRNAs: genomics, evolution, and mechanisms. Cell. 2013; 154(1):26–46.

30. Nott A, Meislin SH, Moore MJ. A quantitative analysis of intron effects on mammalian gene expression. RNA. 2003;9(5):607–17.

31. Shaul O. How introns enhance gene expression. Int J Biochem Cell Biol. 2017;91(Pt B):145–55.

32. Adams B, Dorfler P, Aguzzi A, Kozmik Z, Urbanek P, Maurer-Fogy I, et al. Pax-5 encodes the transcription factor BSAP and is expressed in B lymphocytes, the developing CNS, and adult testis. Genes Dev. 1992;6(9):1589–607.

33. McManus S, Ebert A, Salvagiotto G, Medvedovic J, Sun Q, Tamir I, et al. The transcription factor Pax5 regulates its target genes by recruiting chromatin-modifying proteins in committed B cells. EMBO J. 2011;30(12):2388–404.

34. Gestri G, Osborne RJ, Wyatt AW, Gerrelli D, Gribble S, Stewart H, et al. Reduced TFAP2A function causes variable optic fissure closure and retinal defects and sensitizes eye development to mutations in other morphogenetic regulators. Hum Genet. 2009; 126(6):791–803.

35. Schorle H, Meier P, Buchert M, Jaenisch R, Mitchell PJ. Transcription factor AP-2 essential for cranial closure and craniofacial development. Nature. 1996;381(6579):235–8.

36. Pauls K, Jager R, Weber S, Wardelmann E, Koch A, Buttner R, et al. Transcription factor AP-2gamma, a novel marker of gonocytes and seminomatous germ cell tumors. Int J Cancer. 2005;115(3):470–7.

37. Eckert D, Buhl S, Weber S, Jager R, Schorle H. The AP-2 family of transcription factors. Genome Biol. 2005;6(13):246.

38. Oakley RH, Cidlowski JA. The biology of the glucocorticoid receptor: new signaling mechanisms in health and disease. J Allergy Clin Immunol. 2013;132(5):1033–44.

39. Schultz R, Isola J, Parvinen M, Honkaniemi J, Wikstrom AC, Gustafsson JA, et al. Localization of the glucocorticoid receptor in testis and accessory sexual organs of male rat. Mol Cell Endocrinol. 1993;95(1-2):115–20.

